# Ultra-sensitive detection of *Phytophthora pluvialis* by real-time PCR targeting a mitochondrial gene

**DOI:** 10.1101/2024.12.18.629255

**Authors:** R. O’Neill, E. McLay, L. Nunes Leite, P. Panda, A. Pérez-Sierra, A. Eacock, J. M. Le Boldus, E. A. Stamm, S. Fraser, R. L. McDougal

## Abstract

*Phytophthora pluvialis* is a forest tree pathogen present in the USA, New Zealand, UK and Belgium. Reported hosts include Douglas-fir in the USA, New Zealand, UK and Belgium, as well as tanoak in the USA, radiata pine in New Zealand, Japanese larch and western hemlock in the UK. Disease symptoms range from needle lesions and casting on radiata pine through to twig and stem cankers, and crown dieback on western hemlock, Douglas-fir and Japanese larch. Current detection methods rely on isolation and culture, or PCR using a single-copy gene target with limited sensitivity at low pathogen titre. A qPCR assay targeting a multiple-copy mitochondrial gene was designed to increase sensitivity of *P. pluvialis* detection in forest samples, critical for informing biosecurity, long term disease management and ongoing research. The resulting assay has a detection limit of 12.8 fg mycelial DNA and can detect the pathogen on average 6.12 qPCR cycles before the single-copy target assay. In New Zealand forest samples, the assay was found to consistently detect *P. pluvialis* in all stages of radiata pine needle disease symptoms from early asymptomatic infection through to fully cast needles. The new assay allowed for asymptomatic detection of *P. pluvialis* in radiata pine needle samples four weeks before visual symptoms of disease were observed. The availability of a highly sensitive assay has also enabled rapid and confident diagnostic support of the biosecurity response in the UK during recent detection of *P. pluvialis*. The assay has been used in applications requiring detection at low pathogen titre levels including asymptomatic infection, stream baiting, cast needles and during biosecurity responses, making it a useful tool for effective early detection and management of *P. pluvialis* in affected forests.

## 1. Introduction

*Phytophthora pluvialis* Reeser, Sutton and E Hansen., a pathogen of forest trees, was first described in Oregon, United States of America (USA), during environmental sampling for *Phytophthora* (Reeser 2013). *Phytophthora pluvialis* was initially only rarely found in association with twig and stem cankers on tan oak (*Notholithocarpus densiflorus*) but was later found to cause disease on Douglas-fir (*Pseudotsuga menziesii*) in Oregon (Hansen et al. 2015). Around this time, *P. pluvialis* was also reported as the causal agent of red needle cast (RNC) disease on radiata pine (*Pinus radiata*) and then Douglas-fir in New Zealand (Dick et al. 2014, Hansen et al. 2015), significantly impacting growth of these hosts (Beets et al. 2013, Gomez-Gallego et al. 2019). In 2021, *P. pluvialis* was discovered in the United Kingdom (UK), causing severe decline on western hemlock (*Tsuga heterophylla*) during routine surveillance for *Phytophthora ramorum* (Pérez-Sierra et al. 2022). In the UK, the trees exhibited crown dieback, needle drop, and branch and stem cankers exuding copious resin. Resinous cankers on branches and stems were also observed on Douglas-fir and on Japanese larch (*Larix kaempferi*) (Pérez-Sierra et al. 2024). These latter symptoms and mortality have not been observed in New Zealand on radiata pine, Douglas-fir or western hemlock to date. Mortality of younger western hemlock understory trees was observed, but this is not seen in New Zealand on radiata pine or Douglas-fir. Similarly in Oregon, Washington, and California disease is limited to defoliation of infected Douglas-fir and infection of western hemlock has never been reported. More recently, *P. pluvialis* was also detected in Belgium in watercourses and from Douglas-fir needles (Pirronitto et al. 2024).

The pathogen is thought to originate from the Pacific Northwest of the USA (Brar et al. 2018, Tabima et al. 2021). Populations of *P. pluvialis* in the USA and New Zealand have been compared using both single nucleotide polymorphisms (SNPs) (Brar et al. 2018) and genotype-by-sequencing (Tabima et al. 2021). Both studies demonstrate greater diversity in the USA and the presence of two genotypes of *P. pluvialis* in New Zealand, NZ1 and NZ2, both hypothesised to originate in the USA. The origin of the UK and Belgian populations and their relationship to the USA and New Zealand populations is currently unknown.

The efficient detection of plant pathogens is critical in assisting biosecurity responses, understanding epidemiology, informing disease management strategies and should be of high sensitivity, reliability, and rapidity (Kulik et al. 2020). However, detection can be a challenging task where pathogen biomass endures at low levels within plant tissues or as spores, without visual symptoms (Kulik et al. 2020). The currently available molecular assay for detection of *P. pluvialis* by real-time PCR (qPCR) targets a single copy nuclear gene, ras-related GTP-binding protein 1 (*ypt1*) with primers Ypap2F/2R/P (*ypt1* assay) (McDougal et al. 2021). While single-copy gene targets are useful for identification and quantification of *Phytophthora* species (Ioos et al. 2006, Schena and Cooke 2006, Schena et al. 2008), there are limitations to the sensitivity of detection of a single copy gene for the detection of plant pathogens, especially from infected plant material or during early infection stages. In contrast, assays that amplify multiple copy gene targets have increased sensitivity, greatly increasing detection from plant material (Cooke et al. 2007, Martin et al. 2012). As *P. pluvialis* infection of radiata pine needles is cryptic and sporulation occurs before disease symptom development (Dick et al. 2014), the ability to detect disease early may allow application of control during this asymptomatic stage.

The internal transcribed spacer (ITS) region of nuclear ribosomal DNA is widely recognised as a ‘universal DNA barcode’ marker (Schoch et al. 2012) as it has the highest probability of successful identification over the broadest range of fungi and oomycetes. However, with increasing numbers of identified *Phytophthora* species, the utility of the ITS region as a sensitive diagnostic tool is limited for differentiation of closely related species within clades or sub-clades, leading to limited resolution for diagnostic purposes (Martin et al. 2014). ITS regions have also shown insufficient interspecies variability for reliable determination of oomycete species (Kulik et al. 2020, Foster et al. 2022). Therefore in this study we considered mitochondrial gene (mitogene) targets as an alternative to ITS for diagnostic identification.

The conserved variation in mitogene sequences between *Phytophthora* species has demonstrated utility for diagnostic applications, including highly specific and sensitive qPCR methods and barcoding (Robideau et al. 2011, Bilodeau et al. 2014, Choi et al. 2015, Rojas et al. 2017, Kunadiya et al. 2019). The use of mitochondrial targets for PCR amplification of oomycete DNA is also notably more frequent than in fungi (Kulik et al. 2020), due to their multiple copy nature and noted potential for increased sensitivity in detection (Martin et al. 2014). It has been suggested that efforts to sequence these loci, perhaps in addition to the traditional ITS locus, will increase the availability of data for future evolutionary analyses as well as the development of appropriate diagnostic and monitoring tools (Martin et al. 2014). However, in contrast to barcoding, qPCR assay design requires high species-level variation in DNA across much shorter sequences suitable for the shorter amplicon length required for efficient qPCR amplification or probe annealing. This can be challenging with mitogene sequences where GC content is low and AT homopolymeric runs are common, leading to difficulties in achieving sufficient melting temperature (Tm) for effective primer design (Bustin and Huggett 2017). It is only in recent years with lower costs for high-throughput sequencing that nuclear and mitochondrial genome (mitogenome) sequences for *Phytophthora* species have become available, making this design possible (Srivastava et al. 2020, Kronmiller et al. 2023).

The aim of this study was to develop a diagnostic qPCR assay for *P. pluvialis* detection with improved sensitivity compared to the available single copy *ypt1* nuclear gene target assay (McDougal et al. 2021). Increased sensitivity will enable more accurate detection in samples with low pathogen titre such as asymptomatic infection or spore trap samples, facilitating early detection, effective biosecurity responses, disease management, and research. Here we present a new mitochondrial target qPCR assay for improved sensitivity in detection of *P. pluvialis*. This assay has been instrumental in the characterisation of RNC in New Zealand, as well as assisting in the early response to detection of *P. pluvialis* across forests in the UK during official surveys in 2021-2022.

## 2. Materials and Methods

### 2.1 Isolates and DNA extraction

A total of 57 isolates, including 17 *Phytophthora pluvialis* isolates, 28 isolates of other *Phytophthora* species, and 12 non-*Phytophthora* isolates were used in this study (Table 1 1). DNA from isolates tested at Scion, New Zealand, was extracted during a previous study and stored at -20°C (McDougal et al. 2021). DNA extracts were quantified using a Qubit™ fluorometer (Thermo Fisher Scientific, Waltham, MA, USA) and assessed for quality on a DeNovix DS-11 UV-Vis Spectrophotometer (DeNovix Inc, Wilmington, DE USA). Initial assay screening was carried out with extracts of DNA from *P. pluvialis* NZFS 3608, NZFS 4589 and NZFS 3056 which were quantified, pooled and used as serial dilutions for initial testing and optimisation of annealing temperature and primer concentration. DNA from *P. pluvialis* NZFS 3032 was used for sensitivity testing. All other isolates (Table 1) were used for specificity testing.

**Table 1.**
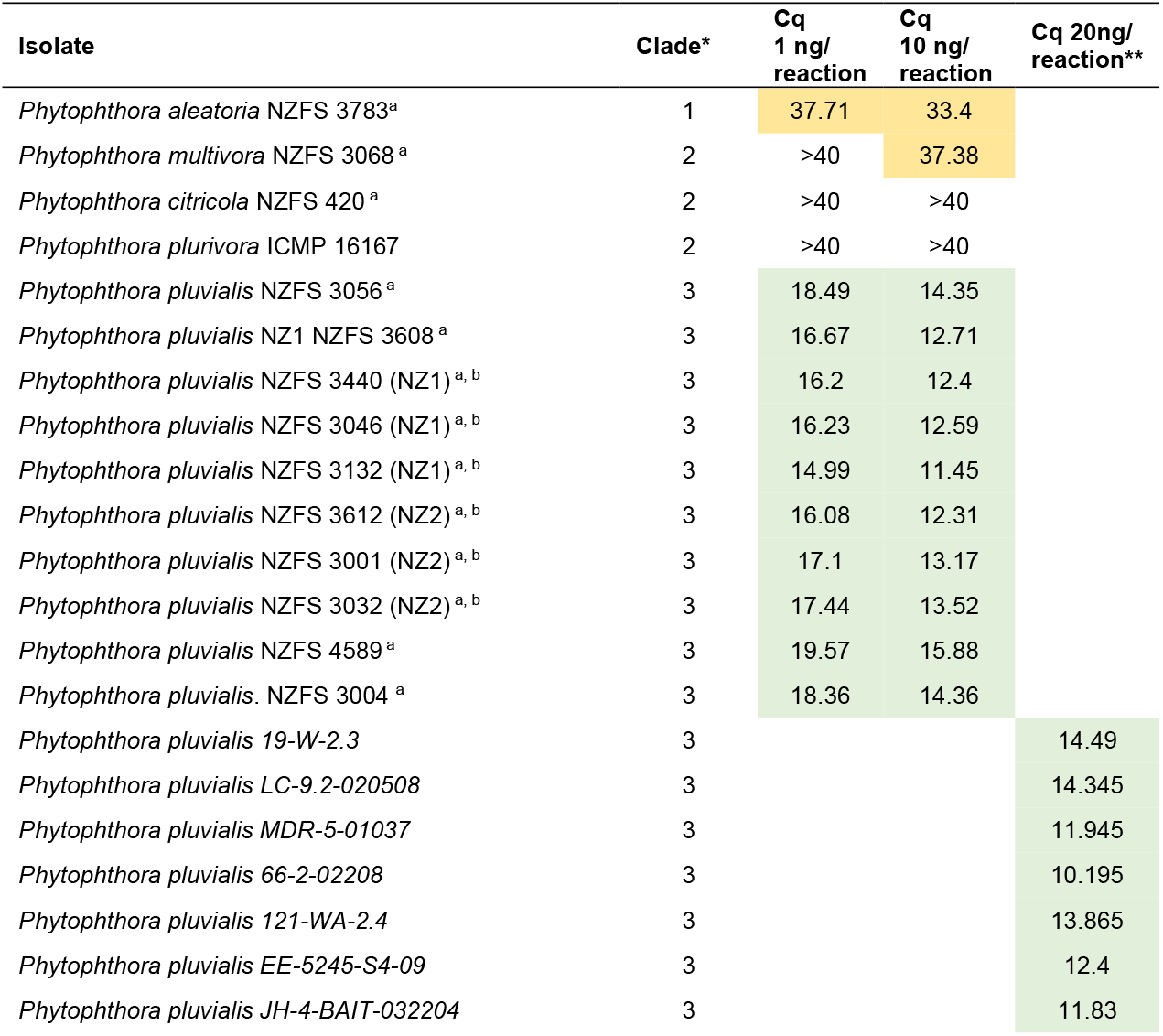
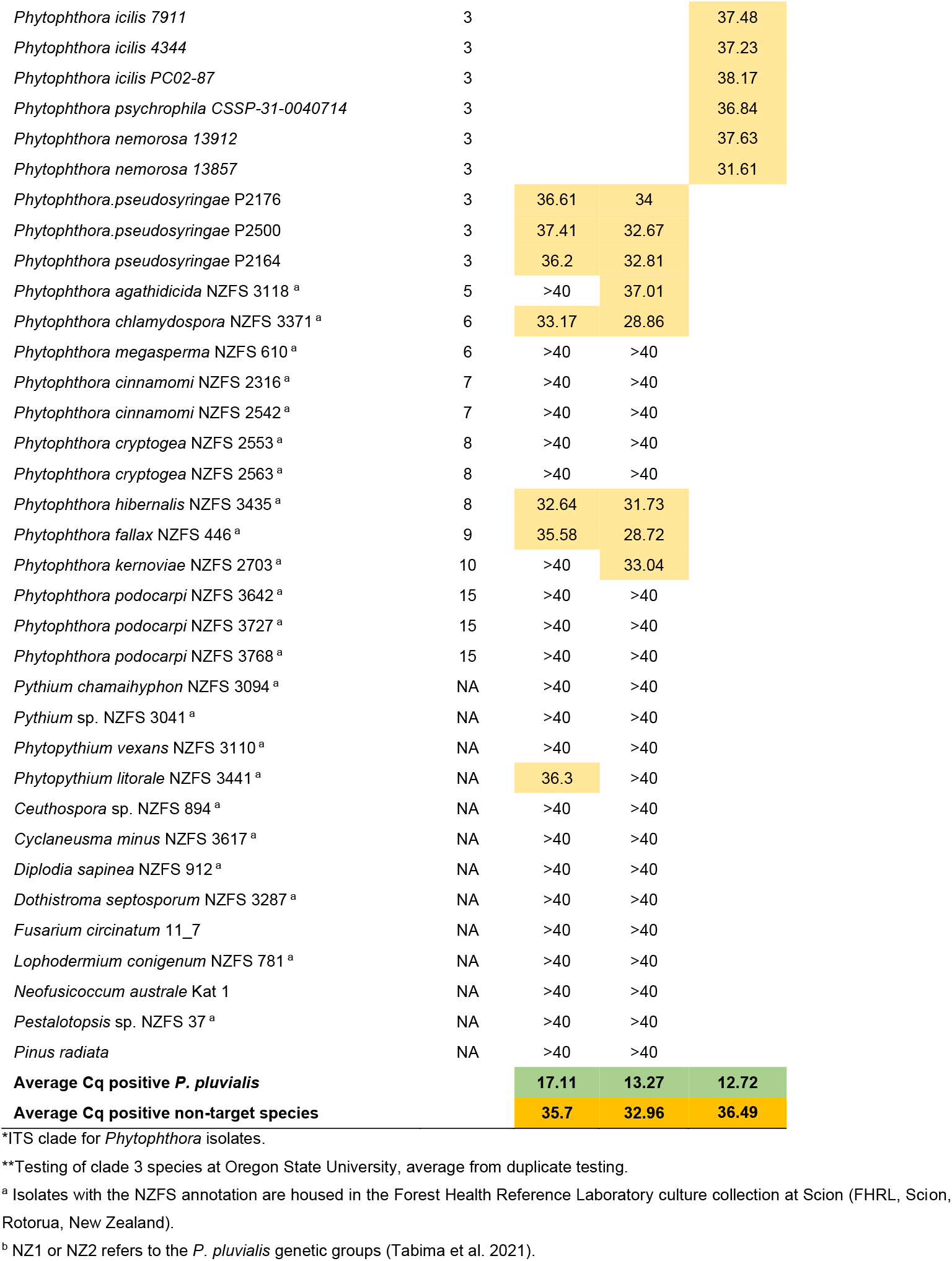
Isolates used in this study and specificity testing results with the *cox2* assay. Results show the Cq for each pure culture DNA concentration. Green coloured cells are those that contain DNA from *P. pluvialis* isolates. Yellow cells are those that gave a Cq value lower than 40 for a non-target species.

Assay sensitivity was also evaluated at Forest Research, UK, using DNA extracted from a pure culture of *P. pluvialis*, TH21/0001, isolated from a branch lesion on western hemlock (*Tsuga heterophylla)* in the UK.

Specificity testing on closely related clade 3 *Phytophthora* species not present in New Zealand (*P. ilicis, P. nemorosa, P. psychrophila*) was carried out in the forest pathology laboratory at Oregon State University (OSU). Isolates were transferred onto V8 media with Whatman Nucleopore Track-Etch 0.4 μm pore membranes placed between the agar plug and the media. After approximately 10 days, mycelium was scraped from the membrane plates and DNA extracted at the Center for Quantitative life Science at Oregon State University (CQLS) using the Mag-Bind Plant DNA DS Kit (Omega Bio-tek, Norcross, GA, United States) with a Kingfisher robot (Thermo Fisher). DNA quality and concentration was assessed using a NanoDrop® ND-1000 UV-Vis Spectrophotometer and Qubit™ fluorometer.

### 2.2 Primer design

Five mitogene targets were identified for qPCR assay design based on mitochondrial phylogenies described in other studies; SecY-independent transporter protein (*ymf16*), 40S ribosomal protein S10 (*rps10*), Cytochrome c oxidase subunit 1 (*cox1*) and Cytochrome c oxidase subunit 2 (*cox2*) (Martin et al. 2014) and NADH dehydrogenase subunit 9 (*nad9*) (Martin et al. 2014, Yuan et al. 2017). Target mitogenes (*ymf16, rps10, nad9, cox1* and *cox2*) were extracted from either annotated *Phytophthora* mitogenomes, or whole genome sequences (Supplementary Table S3) using reference mitogenomes as a custom BLAST database, in Geneious R10 (Kearse et al. 2012). For clade 3 species with no available whole genome sequences, nucleotide sequences of interest from GenBank were added to the primer design alignment (Supplementary Table S3). The resulting DNA reference sequence alignment was used to predict nucleotide differences between *P. pluvialis* and other species for primer design. Particular attention was paid to the closely related clade 3 species *P. ilicis, P. nemorosa, P. pseudosyringae* and *P. psychrophila* which are associated with forests in Oregon and the UK (Kroon et al. 2012, Scanu et al. 2012, Hansen et al. 2017) but are not currently present in New Zealand.

Primer design was carried out with multiple software; Primer 3 Plus (http://www.bioinformatics.nl/cgi-bin/primer3plus/primer3plus.cgi), PrimerBLAST (https://www.ncbi.nlm.nih.gov/tools/primer-blast/) and Geneious R10 to find and cross-check the best candidate primer sets. Primer dimer formation and hairpin formation were evaluated by Oligo Calc (http://biotools.nubic.northwestern.edu/OligoCalc.html). *In-silico* predictive specificity testing against non-target *Phytophthora* and other fungal species was performed using Geneious R10 alignments of primers mapped to *Phytophthora* reference sequences and NCBI PrimerBLAST primer pair specificity parameters using nt database and organism search Oomycetes (taxid:4762).

Due to the low GC content of mitogene target sequences, primer selection criteria were adjusted to less stringent levels: 18 to 30 nt size; melting temperature (Tm) between 58 °C and 64 °C but with Tm difference less than 2 °C between primer pairs, GC content between 45% and 60%, yielding amplicons shorter than 220 bp. To increase specificity, primers were designed where possible to contain mismatches with non-target sequences at the 3′ end.

### 2.3 PCR conditions

End-point PCR was used for gradient annealing temperature optimisation and DNA sequencing to confirm product amplification in this study. PCRs were performed in a total volume of 10 μL, containing 5.4 μL PCR-grade water, 2 μL 5x HOT FIREPol Blend Master Mix 10mM MgCl2 (Solis BioDyne, Tartu, Estonia), 300 nM of each primer and 2 μL template DNA (at both 2 ng/μL and 20 ng/μL). Cycling conditions consisted of an initial denaturing step at 95 °C for 12 min followed by 40 amplification cycles of 15 s at 95 °C, 30 s annealing at 1 °C increments between 54 °C and 61 °C, 1 min at 72 °C and a final elongation for 5 min at 72 °C, run on a SensoQuest Labcycler (SensoQuest GmbH, Göttingen, Germany). PCR products were run on a 1.5% agarose gel (Bioline Reagents LTD, London, UK) in Invitrogen™ UltraPure™ TBE Buffer (Thermo Fisher Scientific, MA, USA) with 2 μL of Invitrogen™ TrackIt™ 1 Kb Plus DNA Ladder at 7 V/cm and stained with RedSafe™ Nucleic Acid Staining Solution (20,000x) (iNtRON Biotechnology, Inc., Seongnam, Republic of Korea) for UV transillumination.

PCR products for sequencing were cleaned up with Exonuclease 1 and Fast Alkaline Phosphatase (Thermo Fisher Scientific, Waltham, MA USA) according to manufacturer’s instructions. Sanger sequencing of end-point PCR products was carried out at Lincoln University, New Zealand on an ABI 3500xL Genetic Analyser, using ABI PRISM® BigDye® Terminator v3.1 Cycle Sequencing Kit (Applied Biosystems, CA, USA).

SYBR-based qPCR was used for primer concentration optimisation, and initial sensitivity screening for all designed primer sets. SYBR-based qPCR assays were performed in a total volume of 20 μL containing 10 uL of PerfeCta SYBR® Green Fastmix (2X) (Quantabio, Beverly, MA, USA), primer (varied concentrations for testing), DNA template 2 μL (4 ng of DNA in total) and the remainder volume with PCR-grade water. Amplification conditions consisted of an initial denaturing step of 95 °C for 5 min, followed by 40 amplification cycles of 5 s at 95 °C, 15 s at annealing temp (varied during assay development) and 10 s at 72 °C. After amplification, a melting curve analysis was performed by heating the reaction mixture at 95 °C for 15 s, cooling to 60 °C for 30 s and incrementally heating again to 95 °C at 0.3 °C per second.

Hydrolysis probe-based qPCR was used for optimisation of probe concentration, final qPCR assay optimisation and specificity testing. qPCR assays were performed in a total volume of 10 μL, containing 5 μL of PerfeCTa qPCR ToughMix® 2X (Quantabio, Beverly, MA, USA), 800 nM of each primer, 200 nM probe, 2 μL of template DNA (4 ng in total) and the remainder with PCR-grade water. Cycling conditions consisted of an initial denaturing step at 95 °C for 5 min followed by 40 amplification cycles of 5 s at 95 °C, 30 s at 58 °C and 10 s at 72 °C.

All PCR runs included a no template control (NTC; PCR-grade water). qPCR analysis carried out at Scion, New Zealand, was carried out on the Mic Real-Time PCR System (Bio Molecular Systems, QLD, Australia), with threshold and quantification cycle (Cq) set automatically by the software.

For testing in Oregon, USA, qPCR was performed on a CFX96 Real-Time System (Bio-Rad, Hercules, CA United States) using SsoAdvanced Universal Probes Supermix (Bio-Rad) under the conditions outlined in Supplementary S1.

### 2.4 Assay optimisation

Temperature gradient end-point PCR was carried out to investigate optimal annealing temperature for each of the primer sets from 54 °C to 61 °C, in 1 °C increments using *P. pluvialis* DNA. Sanger sequencing of end-point PCR products was carried out to confirm amplification of the desired template sequence, with alignment against the target *P. pluvialis* gene regions in Geneious R10, along with BLAST where available (Altschul et al. 1990). qPCR was later carried out using the hydrolysis probes designed for successful candidate assays, to confirm the annealing temperature of the probe-based assay for qPCR.

Primer concentration was assessed by SYBR-based qPCR in a concentration series from 200 nM up to 800 nM per reaction, in 100 nM increments including forward and reverse primer concentration variation. Hydrolysis probe concentration optimisation and sensitivity testing was then carried out with 100, 150, 200, 250, 300 and 350 nM final reaction probe concentrations. A final assessment of optimal annealing temperature including the hydrolysis probe was also carried out. Both two-step and three-step qPCR protocols were investigated to optimise results.

### 2.5 Sensitivity and limits of detection (LOD)

Assay sensitivity was tested using 5-fold serially diluted *P. pluvialis* DNA (2.1) diluted in sterile PCR-grade water to yield 11 concentrations ranging from 25 ng per reaction down to 2.56 fg. This dilution series data was used to generate standard curves, determine the limit of detection (LOD) of the assays, and to compare the sensitivity of mitogene target assays with that of the *ypt1* assay (McDougal et al. 2021). Reactions were run in technical replicates of six using hydrolysis probe-based qPCR (2.3). The assay efficiency, coefficient of determination (R^2^) and y-axis intercept were calculated for each assay as parameters to assess the differences between potential candidates.

Assay sensitivity was also evaluated at Forest Research, UK (2.1). Ten-fold serial dilutions of DNA were made using PCR-grade water to yield concentrations ranging from 1.2 ng down to 1.2 pg per reaction. Reactions were repeated six times in separate runs and performed in technical triplicates in each run, with qPCR conditions as per Supplementary S1. The LOD of the assays was defined as the lowest concentration of DNA for which ≥ 95% of replicates gave positive results (Supplementary Table S7, S8).

### 2.6 Specificity testing

Assay specificity was tested on DNA extracted from pure cultures of a wide variety of species (Table 1). The clade 3 *Phytophthora* species present in New Zealand were assessed using 10 isolates of *P. pluvialis*, and three isolates of *P. pseudosyringae. Phytophthora* clades 1, 2, 5, 6, 7, 8, 9, 10, and 15 were also assessed using a variety of species commonly associated with forest tree samples. Four other oomycetes in genera *Pythium* and *Phytopythium*, eight species of forest pathogenic fungi, and radiata pine were included in the specificity tests. DNA samples were diluted in PCR-grade water to 10 ng/μL and 1ng/μL before testing. Reactions were run in triplicate using hydrolysis probe-based qPCR under the conditions outlined above.

The other clade 3 *Phytophthora* species not present in New Zealand (*P. ilicis, P. nemorosa, P. psychrophila*) were tested for potential assay interaction by the LeBoldus laboratory at Oregon State University. Three *P. ilicis* isolates, two *P. nemorosa* isolates, one *P. psycrophila* isolate and seven *P. pluvialis* isolates were. All DNA concentrations were adjusted to 20 ng/μL in PCR-grade water prior to qPCR according to conditions in Supplementary S1, and tested in duplicate with a NTC per plate. qPCR was performed with a Bio-Rad CFX96 Real-Time system (Bio-Rad, Hercules, California, USA).

### 2.7 Detection of P. pluvialis from infected plant samples

#### 2.7.1 Controlled inoculation of detached needles with *Phytophthora pluvialis* zoospores

Controlled inoculations of radiata pine needles with *P. pluvialis* were used to assess the sensitivity of assay detection in a background of host tree DNA. Radiata pine needles were collected from a Bay of Plenty forest, New Zealand following a negative qPCR test (McDougal et al. 2021) a month prior, and lack of lesions or damage at the time of collection. Detached needles were either stored at 4 °C darkness or infected with *P. pluvialis* (NZFS 4528). Inoculation was performed by placing needles in a 50 mL falcon tube containing 45 mL of zoospore solution (2.25 × 10^4^ spores mL^−1^) for seven days at 17 °C in darkness. Needles were surface sterilised for 30 seconds in 70% ethanol then rinsed twice consecutively with sterile deionised water for 30 seconds. Uninoculated tissue was processed for qPCR and distributed into wells before inoculated needles were removed from the growth chamber, to reduce the risk of cross-contamination from inoculated material. Symptomatic needle tissue was then cut into 1 to 2 mm fragments and weighed along with uninoculated needle tissue to achieve mixtures with varying percentages of inoculated and uninoculated tissue at 0, 1, 10, 25, 50, 75, 100% inoculated tissue. Samples of each inoculation mixture were placed into a 96 deep U-bottom 2 mL well plate (GEB, China) in replicates of 10 and frozen at -20 °C for qPCR testing to investigate the sensitivity of detection from pine host plant material. Metalware used in sample preparation was bleached and flame dried in between each sample. Each of these inoculated mixtures were tested with the *ypt1* assay for *P. pluvialis* detection (McDougal et al. 2021), the *cad* target assay CAD918/CAD1019/Probe945 (*cad* assay) for reference detection of the radiata pine host (Chettri et al. 2012) as well as the candidate assay designed in this study through a high-throughput commercial PCR provider (Slipstream Automation, Palmerston North, New Zealand. Further details supplied in Supplementary materials S2).

#### 2.7.2 qPCR assay evaluation across a variety of typical RNC symptoms in field samples

To assess assay performance in field conditions, needles expressing symptoms typical of RNC following natural infection in a forest within the East Coast of New Zealand’s North Island were tested. At a needle level, RNC infections progress from asymptomatic to the formation of olive-coloured lesions, often containing resin bands. Lesions turn red, then brown over time (Dick et al. 2014) and may be small or can also extend the length of the needle. Needle fascicles were collected from four 17-year-old *Pinus radiata* trees in an east coast forest in the North Island of New Zealand. Needle fascicles were stored at 4 °C in darkness for 2 days. Each needle was scored as a percent of total area displaying symptoms (lesions/resin bands) typical of RNC (Figure 3C). The needles were classified into 7 levels of severity according to the proportion of disease tissue: 0% (asymptomatic), 10% olive lesions, 20% mixed lesions, 30% olive lesions, 50% mixed lesions, 70% brown lesions and 80% red lesions. Samples were prepared for DNA extraction and qPCR analysis as per 2.7.1.

**Figure 1.**
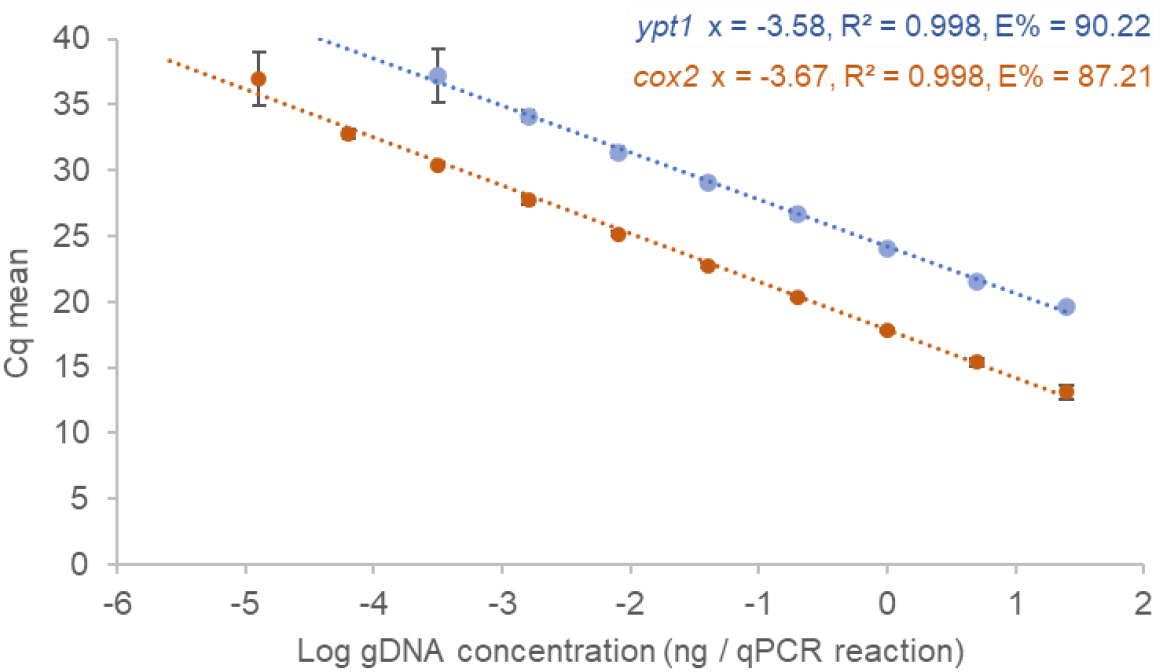
Standard curves for *cox2* and *ypt1* (McDougal et al. 2021) assays using a 5-fold serial dilution of *P. pluvialis* (NZFS 3032) DNA. qPCR was carried out using six technical replicates. Coefficient of determination (R^2^), slope and amplification efficiency including the LOD limits are indicated for each assay (320 fg for the *ypt1* assay and 12.8 fg for the *cox2* assay). Error bars represent the standard deviation (SD). The data point showing the highest Cq value is the LOD for that assay. Neither assay showed positive amplification at the lowest serial DNA dilutions corresponding to 2.56 fg DNA per qPCR reaction.

**Figure 2.**
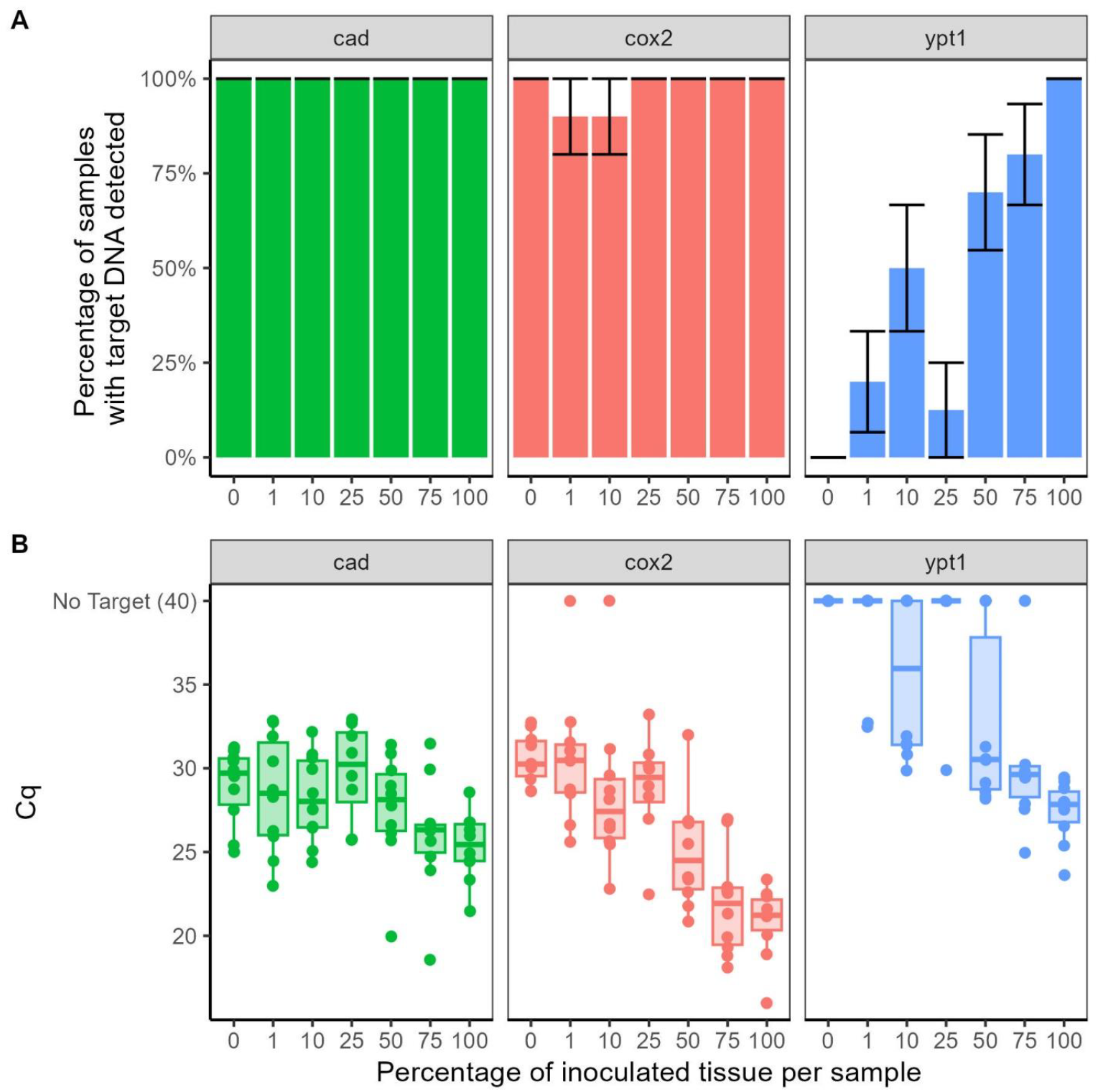
The sensitivity of qPCR assays targeting cox2 (red) and ypt1 (blue) genes for detection of *Phytophthora pluvialis*, and host *Pinus radiata* (cad, green) over increasing proportion of inoculated radiata pine needle tissue. A; Percentage of samples in which target DNA was detected by each assay. Error bars indicate standard error. B; the Cq value of each inoculum mixture where samples with no target detection (“No Target”) are set to a maximum value of 40. Dots indicate individual samples; Boxes indicate median and quartile ranges.

**Figure 3.**
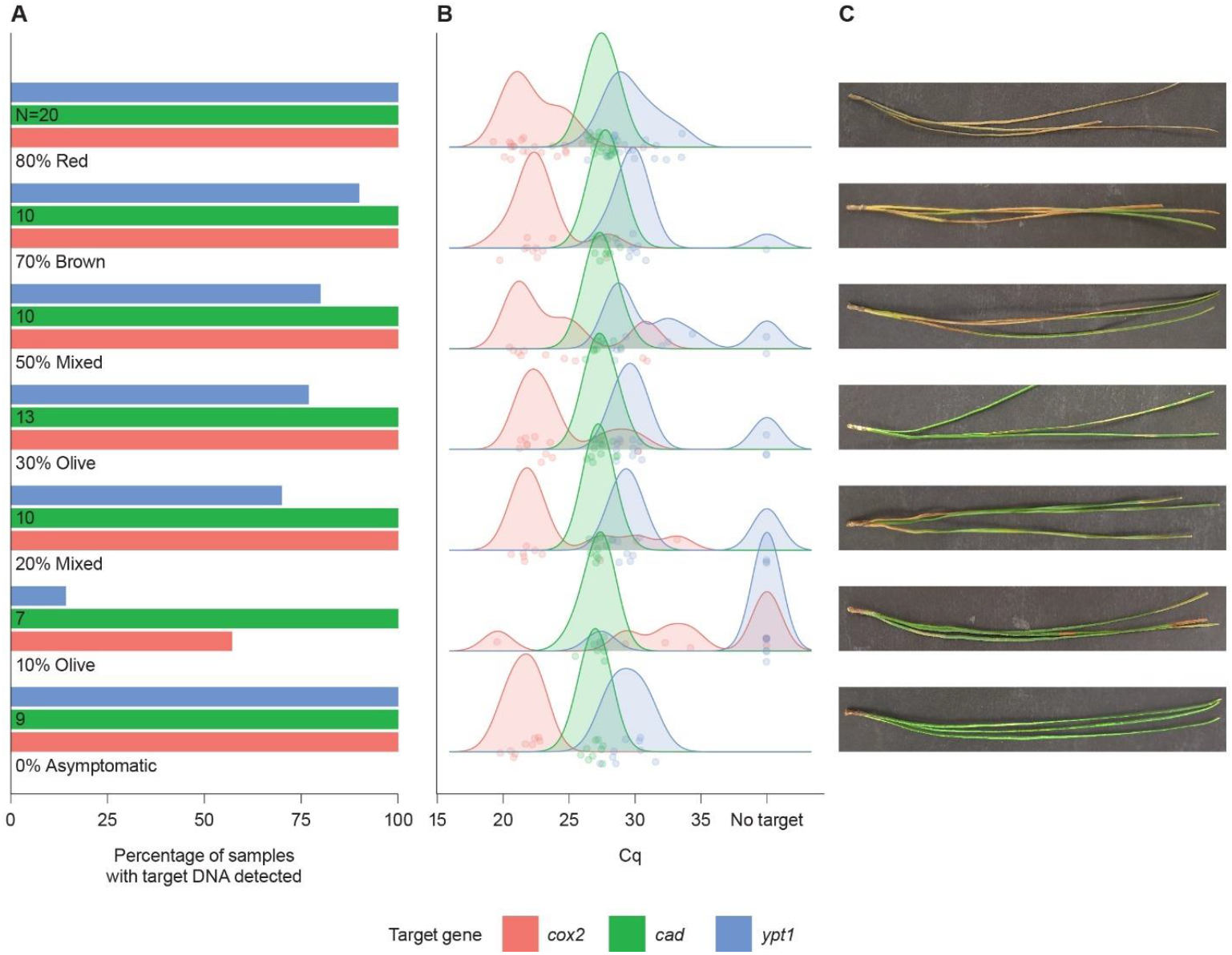
The sensitivity of qPCR assays targeting cox2 (red) and ypt1 (blue) genes for detection of *Phytophthora pluvialis*, and host *Pinus radiata* (cad, green) from samples collected from a New Zealand forest experiencing a natural disease outbreak. Samples were categorised into a range of symptom severities from 0-80+ % severity. A; Percentage of samples in which target gene was detected by each assay. The number of samples tested for each sample type (N) is shown at the top of the bar. B; the Cq value of each sample as a jittered point. A density curve is shown for each target gene to show general trends in performance where non detection is set to a maximum value of 40. C; Example of symptoms within each category.

#### 2.7.3 Routine monitoring of cast needles in New Zealand radiata pine forests

The assay developed in this study was used to analyse needle samples from a central North Island forest in New Zealand. Samples of cast needles were collected regularly from needle traps during 2022 and 2023 regardless of the presence of visual symptoms in overhead canopy. In total, 556 samples were analysed. Needle samples collected from traps were processed for qPCR analysis as per 2.7.1

#### 2.7.4 Testing of infected bark, wood, needles and water baits in the United Kingdom

During 2021-2022, samples of western hemlock and Douglas-fir from official surveys in response to the first detection of *P. pluvialis* in the UK were tested with the new assay. Sample preparation was performed using aseptic methods and flame-sterilised tools. Potentially infected plant tissues, such as needles or bark from tree branches or from western hemlock water baits (needles) were selected. Pieces of approximately 0.5 - 1 cm^2^ were initially collected, then subsequently cut into smaller pieces (maximum 2 × 2 mm, each side). DNA was extracted on the day of receipt, or samples were stored at -20°C until required. A maximum of 50 mg of tissue was processed for DNA extraction using the DNeasy Plant Pro DNA extraction kit (Qiagen, Hilden, Germany). DNA extraction was performed with modifications to the standard protocol to include the addition of 450 μL of CD1 (lysis buffer) and 50 μL of PS solution during lysis to account for high levels of phenolic compounds. Lysis of plant tissue was performed using the PowerLyzer24 (Qiagen, Hilden, Germany) for three cycles of 20 seconds at 2000 RPM, with a dwell time of five minutes between each cycle. Each extraction run contained a negative extraction control (NEC), to check for cross-contamination, and qPCR conditions as per (Supplementary S1).

## 3. Results

### 3.1 Assessment and optimisation of candidate assays

Four of the five target mitogenes assessed in this study were suitable for primer design (*rps10, cox1, cox2* and *nad9)* based on sufficient variation in the reference sequence alignment for differentiation of species. *S*even qPCR assays were designed across these four regions for testing. *In-silico* analyses of all the designed primer sets did not predict primer dimers or secondary structures of concern for PCR, and neither did Geneious alignment or PrimerBLAST analyses reveal potential specificity issues. At each step of assay testing and optimisation, unsuccessful assays were discarded and successful assays progressed.

All designed primer sets produced the predicted size PCR products at every annealing temperature from 54 °C to 61 °C in the temperature gradient end-point PCR. The best temperatures for continued optimisation were chosen based on the highest yield end-point PCR bands visualised on 1.5% agarose gel (data not shown). For the qPCR assay testing, the best annealing temperature was that with the lowest Cq value, assay efficiency closest to 100% and R^2^ value closest to 1. Optimised primer and probe concentrations for qPCR were the concentration producing the lowest Cq, efficiency closest to 100% and R^2^ closest to 1. The *cox2* target primer set cox2_581F/2R/2P (*cox2* assay) was selected as the successful candidate assay based on these and subsequent testing. The primer and probe sequences are cox2_581F (5’-ATGGTTGCCGGAATTTTATGAGTT-3’), cox2_2R (5’-GGCAGAACCTTGGCAATTAGG-3’) and cox2_2PL (5’-FAM-TACCTTCCATAACTGGAGTTGCGGGATCT-BHQ1-3’), with a product size of 177 bp. The optimal primer concentration for cox2-581F/2R/2P (*cox2* assay) is 800 nM for each primer, and 200 nM for the hydrolysis probe. The optimal annealing temperature for this assay is 58 °C. Final hydrolysis probe-based qPCR cycling conditions for this assay are outlined in 2.3.

DNA sequencing of the cox2_581F/2R end-point PCR product yielded confirmation of the correct sequence amplification. When aligning the sequenced PCR product with original primer design reference sequences, the PCR product matched the *P. pluvialis* reference sequence, and the *P. pluvialis* whole genome NCBI accession LC596038.1 with 100% identity. The nearest clade 3 *Phytophthora* species have 97% identity to this sequence and many other species have far less (Supplementary Table S6).

### 3.2 Evaluation of sensitivity, LOD and standard curve generation

The sensitivity was evaluated for the *cox2* assay and compared to the *ypt1* assay (McDougal et al. 2021). The LOD was defined as the lowest amount of DNA template at which at least one replicate was positive (Verdecchia et al. 2021). The LOD for the *cox2* assay was 12.8 fg DNA per reaction, compared to 320 fg for the *ypt1* assay, though this was originally published as 2 pg (2000 fg) on a different qPCR instrument and on different isolates of *P. pluvialis* (McDougal et al. 2021). For the isolate of *P. pluvialis* tested in the UK, the *cox2* assay was able to detect down to 120 pg DNA per reaction. On average, the *cox2* (multiple copy mitogene target) assay detected *P. pluvialis* DNA 6.12 qPCR cycles before the *ypt1* (single copy nuclear gene target) assay, indicating a 93-fold increase in sensitivity (Supplementary Table S7).

### 3.3 Specificity testing

*In-silico* specificity testing showed no non-specific product possibilities in primerBLAST for the *cox2* assay which was the most specific option of all designed candidate assays. Within the primer and probe sequences there are 18 nucleotides of variance along the reference alignment of non-target species. There are ten nucleotides of variance in the forward primer, three mismatches specific to *P. pluvialis*., and three nucleotides of variance in the reverse primer. The probe contains five nucleotides of variance compared to the reference alignment; one mismatch specific to *P. pluvialis* (Supplementary Figure S5).

*In-vitro* specificity testing (Table 1) differentiated *P. pluvialis* DNA from the other species tested. All *P. pluvialis* isolates successfully tested positive, with an average Cq of 13.27 and 17.11 for 10 ng and 1 ng of DNA input per reaction, respectively. Amplification was also observed for other *Phytophthora species*, but at a later amplification cycle, with average Cq of 32.96 and 35.7 for 10 ng and 1 ng of DNA input per reaction. Clade 3 species tested in Oregon at 20 ng of DNA input per reaction showed amplification with an average Cq of 36.49, whereas the *P. pluvialis* isolates were effectively identified with an average Cq of 12.72. Late Cq amplification for other *Phytophthora* isolates outside of clade 3 was observed for *P. aleatoria, P. multivora, P. pseudosyringae, P. agathidicida, P. chlamydospora, P. hibernalis, P. fallax* and *P. kernoviae*. The only late amplification observed outside of the genus *Phytophthora* was for *Phytopythium litorale*, at similar quantification cycles as that for the other non-specific amplification. No amplification was observed for *Phytophthora* isolates belonging to clades 7 and 15, as well for the fungi and the radiata pine host.

### 3.4 Detection of P. pluvialis from infected pine samples

#### 3.4.1 Controlled inoculation of detached needles with *Phytophthora pluvialis zoospores*

*Phytophthora pluvialis* DNA was detected by the *cox2* assay in a greater proportion of samples, with greater sensitivity in most cases than the *ypt1* assay (**Error! Reference source not found**.). High levels of detection of *P. pluvialis* (positive in 90 to 100% of samples) occurred in all samples across the series when tested with the *cox2* assay, including the uninoculated control sample (**Error! Reference source not found**.*A*). In contrast, the proportion of samples with *P. pluvialis* detection increased with the proportion of inoculated host tissue when tested with the *ypt1* assay, except for samples with 25% inoculated tissue, in which only 12.5% of samples tested positive. The Cq values obtained with both assays decreased with increased proportion of inoculated tissue, while the Cq values obtained with the pine assay (*cad*) remained relatively stable (standard error of 0.33) (**Error! Reference source not found**.*B*). The Cq was consistently lower when a sample was tested with the *cox2* assay compared to the *ypt1* assay, indicating a higher sensitivity (Supplementary Table S7). Due to the detection of *P. pluvialis* in the uninoculated (0%) sample, the collection site was revisited four weeks after the initial sampling and mild RNC symptoms were observed on the trees. Samples were again collected, and isolations performed as described by Hood *et al*., ((Hood et al. 2022)). The resulting culture was identified by ITS-6/4 sequencing (White et al. 1990, Cooke et al. 2000) as *P. pluvialis* (NZFS 5557).

#### 3.4.2 Evaluation of the *cox2* assay across a variety of typical red needle cast symptoms in field samples

The *cox2* assay detected *P. pluvialis* in 100% of samples across all sample types except those with small olive lesions (∼ 10% of the needle length) in which only 57% of samples tested positive (**Error! Reference source not found**.A). *Phytophthora pluvialis* detection with the *ypt1* assay was also lowest in samples with 10% olive lesions (14% detection), and detection success increased with lesion size regardless of symptoms type (olive, red, brown or mixed). All asymptomatic samples tested positive for *P. pluvialis* with both assays. The distribution of Cq values was generally lower across samples when tested with the *cox2* assay compared to the *ypt1* assay (**Error! Reference source not found**.B). The Cq for radiata pine (*cad*) remained consistent across samples. The peak Cq for the *cox2* assay was relativity consistent (20 to 25 Cq) for all samples except 10% olive (Figure 3 A/B). However, the peak Cq with the *ypt1* assay was more varied, and generally above 28, otherwise not detected.

#### 3.4.3 Routine monitoring of cast needles in New Zealand

Cast radiata pine needles were collected from needle traps within monitoring plots in central North Island of New Zealand in 2022 and 2023 (n = 556). In total, *P. pluvialis* was detected from 147 samples using the *cox2* assay and 40 samples using the *ypt1* assay. *Phytophthora pluvialis* was detected in a greater number of samples in 2023 than 2022 (Figure 4). A seasonal pattern was observed in both regions with a peak in positive detections in July to September (2022; 42% positive, 2023; 71% positive), and the fewest detections in Jan to March (2022; 0% positive, 2023; 9% positive) using the *cox2* assay within their respective years. A similar trend was observed using the *ypt1* assay, however a reduced number of positives were detected in Jan to March 2023 (1% positive) and July to September of both years (2022; 11% positive, 2023; 25% positive). The Cq values for positively assigned samples ranged between 19.4 to 35.5 using the *cox2* assay and 26.6 to 34.8 using the *ypt1* assay (Figure 4B).

**Figure 4.**
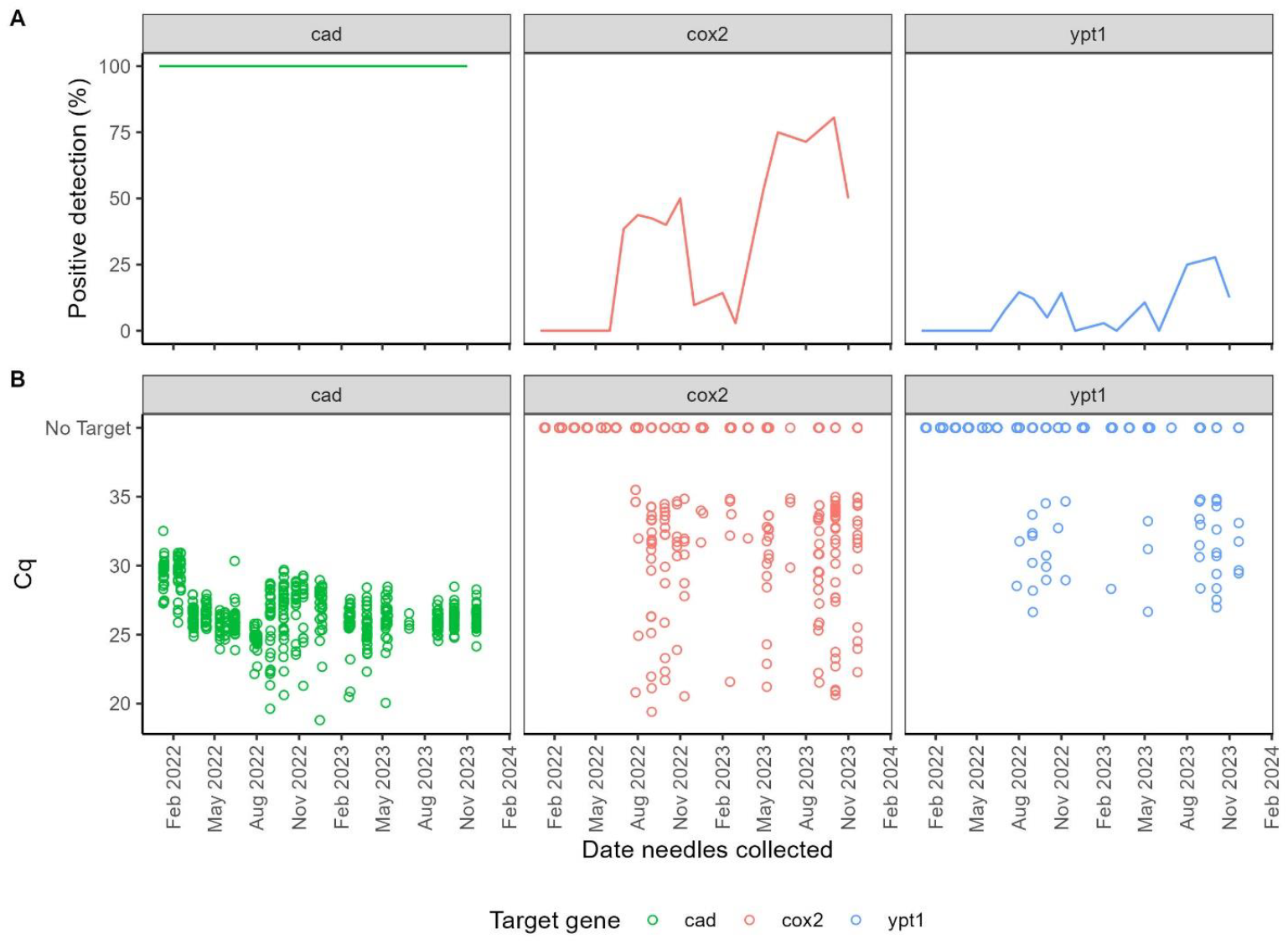
Detection of *Phytophthora pluvialis* using the cox2 (red) or ypt1 (blue) assay with a background host *Pinus radiata* (cad assay, green). A) Percentage of samples per month which tested positive. B) Cq of each sample with negative samples shown as no target.

#### 3.4.4 Testing of infected bark, wood, needles and water baits from forests in the United Kingdom

A total of 1361 samples of western hemlock were tested in 2021-2022. 58% of the samples were collected in England, 17% in Scotland, and 25% in Wales. In total 21% of samples tested positive for *P. pluvialis*, 176 from England, 28 from Scotland and 81 from Wales. Most of the positive samples were obtained from resinous cankers either on branches or stems (n=257), with the others coming from needles (n=19) or needle litter (n=9). The Cq values for positively-assigned samples ranged between 15.37 and 34.6. Western hemlock water baits (needles) were also tested. In total 96 of 161 samples tested positive, with Cq values ranging between 19.12 and 34.6. A total of 351 samples of Douglas-fir were tested, with 11% testing positive, 24 from England, three from Scotland and 10 from Wales. The Cq values ranged between 21.23 and 34.42.

## 4. Discussion

*Phytophthora pluvialis* is a significant pathogen of commercial conifer trees in New Zealand, USA, UK and most recently Belgium (Reeser 2013, Dick et al. 2014, Hansen et al. 2015, Pérez-Sierra et al. 2022, McLay et al. 2023, Pirronitto et al. 2024). Our ability to detect the pathogen early and rapidly is critical for the prevention of spread, long-term disease management, and research activities. The aim of this study was to develop a multiple-copy gene target qPCR for detection of *P. pluvialis* with greater sensitivity than the currently available single-copy *ypt1* target assay Ypap2F/2R/P (McDougal et al. 2021). Targeting mitogenes for qPCR assay design is an effective solution for increased detection sensitivity due to the multiple copy nature of mitochondrial DNA in eukaryotic cells (Kulik et al. 2020, LeBlanc et al. 2022). We designed and tested seven different qPCR primer sets targeting four different mitogenes in *P. pluvialis*. Assessment of optimal annealing temperature, primer concentration, sensitivity and specificity of these primers were used to select the best performing candidate primer set, targeting the *cox2* gene. The cox2_581F/2R/2P *cox2* qPCR assay developed in this study proved to be 93-fold more sensitive in detection of *P. pluvialis* from infected pine needles than the *ypt1* assay.

Based on genomic analyses of the New Zealand and USA populations it was previously thought that *P. pluvialis* had originated from the USA (Brar et al. 2018, Tabima et al. 2021) with a single introduction likely followed by clonal expansion in New Zealand (Tabima et al. 2021). It is suspected that *P. pluvialis* may have been introduced to NZ on imported forest machinery. However, it unknown how *P. pluvialis* arrived in the UK, and more recently Belgium. The use of the NZ1 and NZ2 genetic variants of *P. pluvialis* (Tabima et al. 2021), along with isolates found in the USA and UK, was important to ensure that no difference in detection would be observed. With the expanding range of this pathogen, this new and highly sensitive diagnostic tool will be critical to tracking the spread of *P. pluvialis* and strengthening global biosecurity efforts.

Testing the specificity of the *cox2* assay with DNA from various *Phytophthora* species and other oomycete and fungal species demonstrated that this assay is specific for *P. pluvialis* under the conditions described here. While some potentially positive Cq values were obtained from DNA at 1 ng or higher per reaction from other non-target *Phytophthora* species, Cq values were readily distinguishable from those of target *P. pluvialis* (Table 1). The other closely related clade 3 species that showed this cross-amplification have not currently been identified in New Zealand pine forests (McDougal et al. 2021). Furthermore, clade 3 species *P. ilicis* and *P. psychrophila* have been noted as being rarely detected in forests at all (Hansen et al. 2017). However, in other locations such as USA, UK and Belgium where *P. pluvialis* has been detected, other clade 3 species are present. Some late Cq amplification occurred from non-target *Phytophthora* species outside of clade 3 also. Of particular note for New Zealand radiata pine forests is *P. kernoviae*, which produced a Cq value of 33.04 from DNA of 10ng per reaction, however at 1ng there was no amplification (Table 1). Symptoms of radiata pine needle infection by *P. kernoviae* and *P. pluvialis* are indistinguishable, and both pathogens are described as casual agents of RNC (Dick et al. 2014). *Phytophthora kernoviae* has also been associated with another needle disease of radiata pine, previously called physiological needle blight (McDougal and Ganley 2021). Some non-target cross-reactivity was also observed during development of the *ypt1* assay in a conventional PCR assay and, like the *cox2* assay, this should be considered when employing PCR for diagnostic purposes (McDougal et al. 2021). The expected detection range of the target should also be considered; this *cox2* assay specificity was tested at minimum 1ng DNA from pure mycelial extracts. In practice, at least for environmental samples, 10pg DNA is more likely to represent pathogen presence (Li et al. 2013, Verdecchia et al. 2021) which could be a suitable application for this current assay, though specificity at that level was not tested in this study. While the application of a Cq cut-off value is not always necessary, for this assay we feel it is appropriate to suggest that amplification later than 35 cycles would require additional testing to confirm *P. pluvialis* presence, such as DNA sequencing of PCR amplicons. The *cox2* assay has increased sensitivity benefits for application of qPCR diagnostics in locations that do not have presence of other clade 3 *Phytophthora* species, or for controlled growth-chamber inoculation trials where other potentially cross-reacting species are not present. The assay has also been successfully used in high-throughput scalable robotic platforms which is ideal for this type of diagnostic assay.

Detection of *P. pluvialis* with the *cox2* assay in negative control samples collected from commercial forest for laboratory inoculation trials can be explained by asymptomatic infection. Following high detection (100% of samples) of *P. pluvialis* using the *cox2* assay of uninoculated (0 % inoculated tissue) in the controlled inoculation experiment, the needle sample site (Bay of Plenty, New Zealand) was revisited to investigate (4 weeks post collection). This was confirmed as a true positive detection through observation of RNC symptoms, alongside isolation and confirmation of *P. pluvialis* by DNA sequencing, indicating likely asymptomatic infection of needle material at the initial sample collection. This was further supported with clean no template control (NTC) and extraction control results through robotic testing processes, as well as higher Cq values consistent with lower pathogen titre. The *cox2* assay also detected *P. pluvialis* from asymptomatic needles collected in the east of the North Island in 2023 as part of the natural disease validation. Although the coincidental presence of *P. pluvialis* in uninoculated control samples as well as asymptomatic samples from natural outbreaks may seem unlikely; locating healthy radiata stands, or those which no chemical control applied, was a challenge in 2023 across New Zealand due to an extremely widespread outbreak of foliar disease, particularly in the Gisborne region (east North Island) where over 50% of 2500 plots in a radiata pine forest displayed symptoms severe and widespread enough to be detected with satellite imagery (Watt et al. 2024a). This was also the case in central North Island, as shown by monitoring of cast needles in which greater detection of *P. pluvialis* occurred in all seasons in 2023 compared to 2022 (Table 2). Seasonal patterns observed in needle trap data were similar to those observed by Dick et al., (2014). The variation in detection of *P. pluvialis* in field data removes the concern that the assay will detect *P. pluvialis* ubiquitously across a forest. Although the detection of *P. pluvialis* in asymptomatic needles resulted in no true negative controls in parts of this validation, it also highlighted the possible application of the *cox2* assay for pre-visual or asymptomatic disease detection, which was not previously possible with the *ypt1* assay. The *cox2* assay was also found to consistently detect *P. pluvialis* at a lower Cq than the *ypt1* assay for all needle disease symptom stages in samples from a natural RNC outbreak. Unexpectedly, detection of *P. pluvialis* was lower for the 10% olive samples than asymptomatic samples. Entire needles were diced, however only 200 to 300 μL volume of 10% olive tissue was loaded into each well for extraction and some wells may have received only non-infected sections. The 10% olive samples were taken from a different tree than the asymptomatic needles, and these results may indicate that symptoms are not always representative of level of infection. These results could be explained by a higher uniformity of early-stage infection in the asymptomatic samples, with a lower uniformity of more established pathogen in the 10% olive samples.

Early detection, especially of asymptomatic tissue, is particularly useful for management of RNC of radiata pine. Sporulation of *P. pluvialis* on radiata pine foliage typically occurs on green tissue, in advance of symptom formation (Dick et al. 2014). Under periods of prolonged wetness and cool temperatures (∼10°C), sporulation can begin at 3 days and can occur for at least 14 days in the absence of symptoms (McLay, unpublished data). Under favourable conditions, the entire infection cycle can occur between asymptomatic needles, without detection. The use of a highly sensitive assay which can detect *P. pluvialis* in these situations is highly beneficial for both biosecurity-based diagnostics and for research that aims to improve disease management. Application of qPCR may seem impractical in large scale operations such as plantation forestry, however the use of this assay in combination with other tools, such as a predictive epidemiological model (Watt et al. 2024b) can overcome these challenges of scale. For example, the model may be able to predict areas in which favourable conditions for asymptomatic pathogen spread is occurring and therefore where monitoring should focus. Integration of the assay with stream baiting analysis at key catchments can aid monitoring at large scale in a relatively cost-effective manner.

The availability of this highly sensitive diagnostic assay enabled rapid and confident diagnostics during the response to the recent detection of *P. pluvialis* in the UK (Pérez-Sierra et al. 2022). We note that data is now being generated (data not shown) with the use of this assay for detection of *P. pluvialis* in environmental samples (water filter membranes and rain traps) as well as cambium and bark from tree species affected with cankers, in addition to foliar samples.

## 5. Conclusion

Despite the challenges of primer design targeting *Phytophthora* mitogenes, diversity in the *cox2* region was sufficient to allow design of a sensitive, multiple-copy gene target assay for detection of *P. pluvialis*. The low-titre detection range of the cox2_581F/2R/2P *cox2* assay developed in this study demonstrates much greater sensitivity than the previously available diagnostic assay for *P. pluvialis*. The *cox2* assay was successful in detecting *P. pluvialis* in naturally infected and laboratory-inoculated radiata pine needles in New Zealand, as well as other forest host types in the UK, and presents a valuable tool for detection of asymptomatic infection. These results demonstrate that the *cox2* assay, combined with an appropriate internal reference control assay (Ioos et al. 2006, Chettri et al. 2012) has useful application for early disease detection informing biosecurity responses, treatment and management plans, as well as further research and epidemiological study across different forest host species.

## 6. Acknowledgements

This work was completed as part of the Resilient Forests Programme funded by the Forest Growers Levy Trust and the Ministry for Business, Innovation and Employment through Scion’s Strategic Science Investment Funding. Work at Oregon State University was supported by Grant Number: 2024-67013-42452 (USDA-NIFA) and cooperative agreement number 58-2072-4-024 (USDA-ARS) and 20-PA-11062765-712 (USDA-FS) to JL. Scion’s National Forest Culture Collection (NZFS) housed and maintained the New Zealand isolates used in this study, with added thanks to Forestry Commission England, Scottish Forestry, Natural Resources Wales and the Forest Research Tree Health Diagnostic and Advisory Service (THDAS) for provision of samples for testing in the UK. The authors wish to thank Liz Cunningham and Catherine Banham for technical assistance, Anthony Thrush and John Mackay (dnature diagnostics & research Ltd) for advice on qPCR probe design. Slipstream Automation, Palmerston North are thanked for automated qPCR services and Norma Merrick at Lincoln University for Sanger sequencing services. David Lane, Diya Sen are also thanked for helpful discussion in the development of the diagnostic tool and publication. Dale Corbett is thanked for his help in the production of figure 3. Thanks also to Peter Clinton and Andrew Cridge for manuscript review.

## Supplementary data

Supplementary data to this article is supplied as a separate file: “SUPP data for O’Neill et al. Ultra-sensitive detection of *Phytophthora pluvialis* by real-time PCR”.

## Notes

### Competing Interest Statement

The authors have declared no competing interest.

