## Supplementary Data for "Ultra-sensitive detection of *Phytophthora pluvialis* by real-time PCR targeting a mitochondrial gene"

**S1.**

**qPCR methods and reagents, Forest Research UK**

The qPCR mix consisted of 2X Takyon™ No ROX Probe 2X MasterMix dTTP (Eurogentec, Seraing, Belgium), 800 nM of each forward and reverse primer, 400 nM µM species-specific probe, 2 µL template DNA, made up with molecular grade water (Integrated DNA Technologies, Coralville, IA) to a final volume of 10 µL. Assays were performed in technical triplicates unless otherwise specified with a no template control (NTC) and positive template controls, under the following cycling conditions: 1 cycle of 95 °C for 3 minutes (initial denaturation), 40 cycles of 95 °C for 10 seconds (denaturation), 59 °C for 15 seconds (annealing), 72 °C for 10 seconds (extension). qPCR results were analysed by threshold cycle (Ct) value using the ‘Absolute quantification/ 2nd Derivative Max’ algorithm in the Roche LightCycler 480 ii software (v.1.5.0.39).

**qPCR methods and reagents, Oregon State University, USA**

The qPCR mix consisted of Biorad SsoAdvanced Universal Probes Supermix (Bio-Rad, Hercules, California, USA), 800 nM of each forward and reverse primer, 400 nM species-specific probe, 2 µL template DNA, made up with molecular grade water to a final volume of 10 µL. Assays were performed in technical triplicates unless otherwise specified with a no template control (NTC) and positive template controls, under the following cycling conditions: 1 cycle of 95 °C for 5 minutes (initial denaturation), 40 cycles of 95 °C for 5 seconds (denaturation), 59 °C for 15 seconds (annealing), 72 °C for 10 seconds (extension). qPCR results were analysed by threshold cycle (Ct) value using the Bio-Rad CFX manager 3.1. Cycle thresholds were determined using the single threshold setting.

**S2. Slipstream Automation methods for DNA extraction and qPCR**

DNA extractions and qPCR assays were performed by commercial service provider Slipstream Automation (Palmerston North) using a proprietary CTAB/Chloroform-based extraction method in a high-throughput automated robotic workflow. Assays were performed in simplex 10 μL reaction volumes containing 1X TaqMan Universal Master Mix, 1 μL DNA template, and primer/probe concentrations as reported in 3.2 or as published (Ypap2F/2R/P assay for *P. pluvialis* (McDougal, Cunningham et al. 2021), and the CAD918/CAD1019/Probe945 assay for radiata pine host (Chettri, Calvo et al. 2012)). Reaction conditions included optimised thermal cycling conditions on Slipstream Automation equipment, using the optimal annealing temperature for each assay as published. Two wells of blank extraction controls were included for every 96-well extraction plate, and no template control (NTC) included for every 96-well qPCR run.

**Table S3. Source of DNA sequences extracted for P. pluvialis primer design alignment.**

| **Clade**^a^ | **Species** | **Isolate** | **Sequence source** | **GenBank Accession** | **Reference** |
| --- | --- | --- | --- | --- | --- |
| 1a | *P. cactorum* | LV007 | Whole genome | [NBIJ00000000](https://www.ncbi.nlm.nih.gov/nuccore/NBIJ00000000.1) | (Grenville-Briggs, Kushwaha et al. 2017) |
| 1c | *P. infestans* | 80029 | Mitogenome | [AY894835](https://www.ncbi.nlm.nih.gov/nuccore/AY894835) | (Avila-Adame, Gomez-Alpizar et al. 2006) |
| 1c | *P. andina* | EC3425 | Mitogenome | [HM590419](https://www.ncbi.nlm.nih.gov/nuccore/HM590419.1/) | Unpublished |
| 1c | *P. mirabilis* | PIC99114 | Mitogenome | [HM590421](https://www.ncbi.nlm.nih.gov/nuccore/HM590421.1/) | Unpublished |
| 1c | *P. ipomoae* | PIC99167 | Mitogenome | [HM590420](https://www.ncbi.nlm.nih.gov/nuccore/HM590420.1/) | Unpublished |
| 1c | *P. phaseoli* | PHY P18 | Mitogenome | [HM590418](https://www.ncbi.nlm.nih.gov/nuccore/HM590418) | Unpublished |
| 1d | *P. nicotianae* | n/a | Mitogenome | [NC_035725](https://www.ncbi.nlm.nih.gov/nuccore/NC_035725) | n/a |
| 2 | *P. multivora* | NZFS 3378 | Whole genome | [LGSM00000000](https://www.ncbi.nlm.nih.gov/nuccore/LGSM00000000) | (Studholme, McDougal et al. 2016) |
| 3 | *P. pluvialis* | NZFS 3000 | Whole genome | [LGTT00000000](https://www.ncbi.nlm.nih.gov/nuccore/LGTT00000000) | (Studholme, McDougal et al. 2016) |
| 3 | *P. pseudosyringae* | n/a | Mitogenome | [CM022727](https://www.ncbi.nlm.nih.gov/nuccore/CM022727.1/) | (McGowan, O'Hanlon et al. 2020) |
| 3 | *P. psychrophila^#^* | P10433 | Nucleotide NCBI | [GU222122](https://www.ncbi.nlm.nih.gov/nuccore/GU222122.1/), [HQ261403](https://www.ncbi.nlm.nih.gov/nuccore/HQ261403.1), [JF772005](https://www.ncbi.nlm.nih.gov/nuccore/JF772005.1), [JF770821](https://www.ncbi.nlm.nih.gov/nuccore/JF770821.1) | n/a |
| 3 | *P. ilicis^#^* | P3939/344 | Nucleotide NCBI | [JF770703](https://www.ncbi.nlm.nih.gov/nuccore/JF770703.1/), [JF771863](https://www.ncbi.nlm.nih.gov/nuccore/JF771863.1), [AY129172](https://www.ncbi.nlm.nih.gov/nuccore/AY129172.1), GU222033 | n/a |
| 3 | *P. nemorosa^#^* | P10288,  P16352 | Nucleotide NCBI | [JF770783](https://www.ncbi.nlm.nih.gov/nuccore/JF770783.1), [JF771957](https://www.ncbi.nlm.nih.gov/nuccore/JF771957.1), [HQ261374](https://www.ncbi.nlm.nih.gov/nuccore/HQ261374.1), [GU222084](https://www.ncbi.nlm.nih.gov/nuccore/GU222084.1) | NCBI, (Robideau, De Cock et al. 2011) |
| 5 | *P. agathidicida* | NZFS 3772 | Whole genome | [LGTR00000000](https://www.ncbi.nlm.nih.gov/nuccore/LGTR00000000) | (Studholme, McDougal et al. 2016) |
| 7c | *P. cinnamomi* | NZFS 3750 | Whole genome | [LGSK00000000](https://www.ncbi.nlm.nih.gov/nuccore/LGSK00000000) | (Studholme, McDougal et al. 2016) |
| 7b | *P. sojae* | P6497 | Mitogenome | [DQ832717](https://www.ncbi.nlm.nih.gov/nuccore/DQ832717.1/) | (Martin, Bensasson et al. 2007) |
| 8a | *P. cryptogea* | NZFS 3771 | Mitogenome | Unpublished | Unpublished |
| 8a | *P. sansomeana* | n/a | Mitogenome | [MH936679](https://www.ncbi.nlm.nih.gov/nuccore/MH936679.1/) | n/a |
| 8c | *P. ramorum* | Pr-102 | Mitogenome | [DQ832718](https://www.ncbi.nlm.nih.gov/nuccore/DQ832718.1/) | (Martin, Bensasson et al. 2007) |
| 9b | *P. polonica* | n/a | Mitogenome | [KT946598](https://www.ncbi.nlm.nih.gov/nuccore/KT946598.1/) | n/a |
| 10a | *P. kernoviae* | NZFS 2646 | Whole genome | [JPWV00000000](https://www.ncbi.nlm.nih.gov/nuccore/JPWV00000000) | (Studholme, McDougal et al. 2016) |
| 15 | *P. podocarpi* | NZFS 3727 | Whole genome | [LGSO00000000](https://www.ncbi.nlm.nih.gov/nuccore/LGSO00000000) | (Studholme, McDougal et al. 2016), (Dobbie, Scott et al. 2022) |

^a^ Designation of phylogenetic clade based on (Abad, Burgess et al. 2023)

*^#^* At the time of primer design, the clade 3 *Phytophthora* species had no available whole genome or mitogenome sequences available in GenBank, instead nucleotide sequences were downloaded for each gene region.

**Table S4. Summary of details for *cox2* assay**

| Oligo | Sequence 5’ – 3’ | Optimised assay concentration | Ta | Product size (nt) |
| --- | --- | --- | --- | --- |
| cox2_581F | 5’-ATGGTTGCCGGAATTTTATGAGTT-3’ | 800nM | 58°C | 177 |
| cox2_2R | 5’-GGCAGAACCTTGGCAATTAGG-3’ | 800nM |  |  |
| cox2_2PL | 5’-FAM-TACCTTCCATAACTGGAGTTGCGGGATCT-BHQ1-3’ | 200nM |  |  |

**Figure S5. Primer design alignment for *P. pluvialis* with reference sequences**

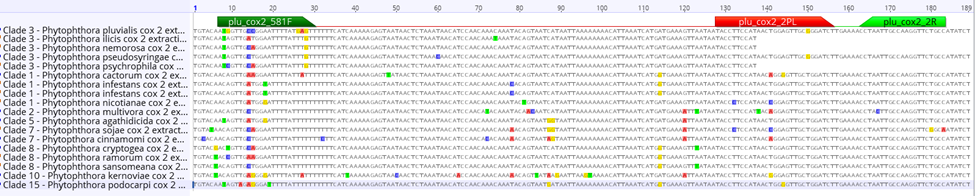

**Table S6. Distance matrix (% identity) between sequenced cox2 PCR products (177 nt) and reference sequences**

| ***Phytophthora* species** | **Accession number** | **Clade** | **Percentage (%) identity with cox2_581F/2R/2P** |
| --- | --- | --- | --- |
| *Phytophthora pluvialis* | LC596038 | 3 | 100 |
| *Phytophthora pluvialis* | LGTT01000265.1 | 3 | 100 |
| *Phytophthora psychrophila* | GU222122 | 3 | 97.66 |
| *Phytophthora pseudosyringae* | CM022727.1 | 3 | 97.44 |
| *Phytophthora ilicis* | GU222033 | 3 | 96.09 |
| *Phytophthora nemorosa* | GU222084 | 3 | 97.66 |
| *Phytophthora cactorum* | NBIJ01000392.1 | 1 | 94.23 |
| *Phytophthora infestans* | AY894835 | 1 | 94.23 |
| *Phytophthora infestans* | NC_002387 | 1 | 94.23 |
| *Phytophthora nicotianae* | NC_035725 | 1 | 93.59 |
| *Phytophthora multivora* | LGSM01000310.1 | 2 | 92.31 |
| *Phytophthora agathidicida* | LGTR01000136.1 | 5 | 93.59 |
| *Phytophthora sojae* | DQ832717 | 7 | 92.95 |
| *Phytophthora cinnamomi* | LGSJ01000489.1 | 7 | 92.31 |
| *Phytophthora ramorum* | DQ832718 | 8 | 95.51 |
| *Phytophthora sansomeana* | MH936679 | 8 | 95.51 |
| *Phytophthora kernoviae* | JPWV02000257.1 | 10 | 89.1 |
| *Phytophthora podocarpi* | LGSN01000280.1 | 15 | 92.95 |

**Table S7. Standard curve data for the *cox2* assay, along with comparative data for the *ypt1* assay, across six* technical replicates using NZFS 3032 (NZ2 variant) *Phytophthora pluvialis* DNA**

| **DNA concentration** | **Assay** | **Cq average** | **Cq Std Dev** | **Efficiency avg %** | **R² average** | **n=*** | **Number cycles difference *ypt1 – cox2*** |
| --- | --- | --- | --- | --- | --- | --- | --- |
| 25ng | cox2_581F/2R/2P | 13.61 | 0.53 | 103 | 0.99 | n=6 | 5.97 |
|  | Ypap2F/2R/P | 19.58 | 0.16 | 87 | 1.00 | n=6 |  |
| 5ng | cox2_581F/2R/2P | 15.42 | 0.29 | 107 | 1.00 | n=6 | 6.15 |
|  | Ypap2F/2R/P | 21.57 | 0.07 | 90 | 1.00 | n=6 |  |
| 1ng | cox2_581F/2R/2P | 17.83 | 0.17 | 102 | 1.00 | n=6 | 6.25 |
|  | Ypap2F/2R/P | 24.08 | 0.15 | 98 | 1.00 | n=6 |  |
| 200pg | cox2_581F/2R/2P | 20.38 | 0.14 | 100 | 1.00 | n=6 | 6.31 |
|  | Ypap2F/2R/P | 26.69 | 0.34 | 100 | 1.00 | n=6 |  |
| 40pg | cox2_581F/2R/2P | 22.78 | 0.13 | 101 | 1.00 | n=6 | 6.30 |
|  | Ypap2F/2R/P | 29.08 | 0.17 | 97 | 1.00 | n=6 |  |
| 8pg | cox2_581F/2R/2P | 25.15 | 0.20 | 100 | 1.00 | n=6 | 6.21 |
|  | Ypap2F/2R/P | 31.36 | 0.40 | 93 | 1.00 | n=6 |  |
| 1.6pg | cox2_581F/2R/2P | 27.71 | 0.34 | 98 | 1.00 | n=6 | 6.44 |
|  | Ypap2F/2R/P | 34.14 | 0.46 | 90 | 1.00 | n=6 |  |
| 320fg | cox2_581F/2R/2P | 30.39 | 0.18 | 99 | 1.00 | n=6 | 5.31 |
|  | Ypap2F/2R/P | 37.17 | 2.03 | 76 | 0.99 | n=2 |  |
| 64fg | cox2_581F/2R/2P | 32.78 | 0.39 | 112 | 1.00 | n=6 | N/A |
|  | Ypap2F/2R/P | - | - | - | - | n=0 |  |
| 12.8fg | cox2_581F/2R/2P | 36.99 | 2.03 | 110 | 0.99 | n=2 | N/A |
|  | Ypap2F/2R/P | - | - | - | - | n=0 |  |
| 2.56fg | cox2_581F/2R/2P | - | - | - | - | n=0 | N/A |
|  | Ypap2F/2R/P | - | - | - | - | n=0 |  |

*This is the number of successful qPCR amplification wells out of the six technical replicates

**Table S8. Additional specificity testing results from Forest Research UK**

| **Isolate** | **Clade** | **Concentration (ng/reaction)** | **Cq** |
| --- | --- | --- | --- |
| *Phytophthora ilicis E02021* | 3 | 30.5 | **35** |
| *Phytophthora ilicis E27821* | 3 | 347.0 | **35** |
| *Phytophthora ilicis E15120* | 3 | 11.5 | **35** |
| *Phytophthora pseudosyringae E37621* | 3 | 399.7 | **35** |
| *Phytophthora pseudosyringae TH21/0329* | 3 | 68.6 | **35** |
| *Phytophthora pseudosyringae 2021/1887* | 3 | 20.1 | **35** |
